# Integrating short-read and long-read single-cell RNA sequencing for comprehensive transcriptome profiling in mouse retina

**DOI:** 10.1101/2024.02.20.581234

**Authors:** Meng Wang, Yumei Li, Jun Wang, Soo Hwan Oh, Rui Chen

**Affiliations:** Department of Molecular and Human Genetics, Baylor College of Medicine, Houston, TX; Human Genome Sequencing Center, Baylor College of Medicine, Houston, TX

**Author notes:** **Corresponding Author** Rui Chen.

**Keywords:** LRS Special Issue, alternative splicing, transcript isoform, long-read sequencing, single-cell RNA sequencing, retinal biology

## Abstract

The vast majority of protein-coding genes in the human genome produce multiple mRNA isoforms through alternative splicing, significantly enhancing the complexity of the transcriptome and proteome. To establish an efficient method for characterizing transcript isoforms within tissue samples, we conducted a systematic comparison between single-cell long-read and conventional short-read RNA sequencing techniques. The transcriptome of approximately 30,000 mouse retina cells was profiled using 1.54 billion Illumina short reads and 1.40 billion Oxford Nanopore long reads. Consequently, we identified 44,325 transcript isoforms, with a notable 38% previously uncharacterized and 17% expressed exclusively in distinct cellular subclasses. We observed that long-read sequencing not only matched the gene expression and cell-type annotation performance of short-read sequencing but also excelled in the precise identification of transcript isoforms. While transcript isoforms are often shared across various cell types, their relative abundance shows considerable cell-type-specific variation. The data generated from our study significantly enhance the existing repertoire of transcript isoforms, thereby establishing a foundational resource for future research into the mechanisms and implications of alternative splicing within retinal biology and its links to related diseases.

## Introduction

The mouse retina is a complex neuronal tissue composed of over 130 unique neuronal cell types that are categorized into seven major cell classes (Masland 2012; Yan et al. 2020; Grünert and Martin 2020). Within each major class, cells can be further classified into sub-classes and cell types. Each cell type differs in its morphology, function, location, and transcriptomic profile (Grünert and Martin 2020; Jeon et al. 1998). Alternative splicing of pre-mRNA is a crucial mechanism that enhances the diversity of transcriptome and proteome (WANG et al. 2015). It plays a pivotal role in cellular differentiation and the development of organisms. Similar to other neural tissues, the retina exhibits a notable enrichment of tissue-specific splicing events (Liu and Zack 2013; Aísa-Marín et al. 2021).

Previous research endeavors have unveiled various aspects of retina-specific splicing, including the identification of retina-specific exons, transcript isoforms, and splicing regulators (Ciampi et al. 2022; Murphy et al. 2016). A comprehensive understanding of the expressed transcript isoforms, coupled with insights into retina cell type-specific splicing events as well as the expression patterns of transcript isoforms at the single-cell level is indispensable for understanding the underlying mechanisms of splicing and gene regulation. Furthermore, a complete catalog of splicing isoforms in individual cell type contexts could guide the accurate prediction of the effect of genetic variants in disorders related to the retina (Aísa-Marín et al. 2021).

Single-cell RNA-sequencing (scRNA-seq) has widely been used to characterize cell-specific transcriptomic differences in various neuronal tissues (Tian et al. 2021). Currently, scRNA-seq data primarily consists of short-read sequencing technologies because of their high accuracy, efficiency, and low cost (Amarasinghe et al. 2020). However, short-read RNA technology is limited to sequencing either the 5’ or 3’ end of the transcript, limiting the ability to quantify RNA transcript isoforms (Byrne et al. 2019). Long-read sequencing is an ideal technology to effectively identify alternative splicing and sequence heterogeneity in mRNA transcripts (Kovaka et al. 2019). Indeed, recent studies of long read-based RNA sequencing (RNA-seq) technology have shown that distinct mRNA transcript isoforms have been observed among different cell types (Tian et al. 2021). Unfortunately, to date, RNA transcript isoforms in the mouse retina have not been systematically annotated and quantified in a cell type-specific context.

In this study, we adapted protocols from the 10x Genomics and implemented a modified pipeline (**Fig. 1A** and **Methods**) to conduct short- and long-read sequencing and data analysis at the single-cell level. The transcriptome of about 30,000 single cells from mouse retinas was profiled with 1.54 billion Illumina short reads and 1.40 billion Oxford Nanopore long reads and a thorough comparison of the short- and long-read single cell RNA-seq technology was performed. Around 17,000 novel isoforms, a 70% increase from previous annotated isoforms, and over 7,000 cell-class specific isoforms were identified. Through differential transcript usage analysis, we found that although transcript isoforms were expressed across many cell types, their usage was varied. Surprisingly, over 1,000 fusion transcripts have been identified, with various interesting features. Taken together, our study generated a comprehensive atlas of full-length transcript isoforms in the mouse retina at the single-cell level, providing an invaluable resource for the community.

**Figure 1.**
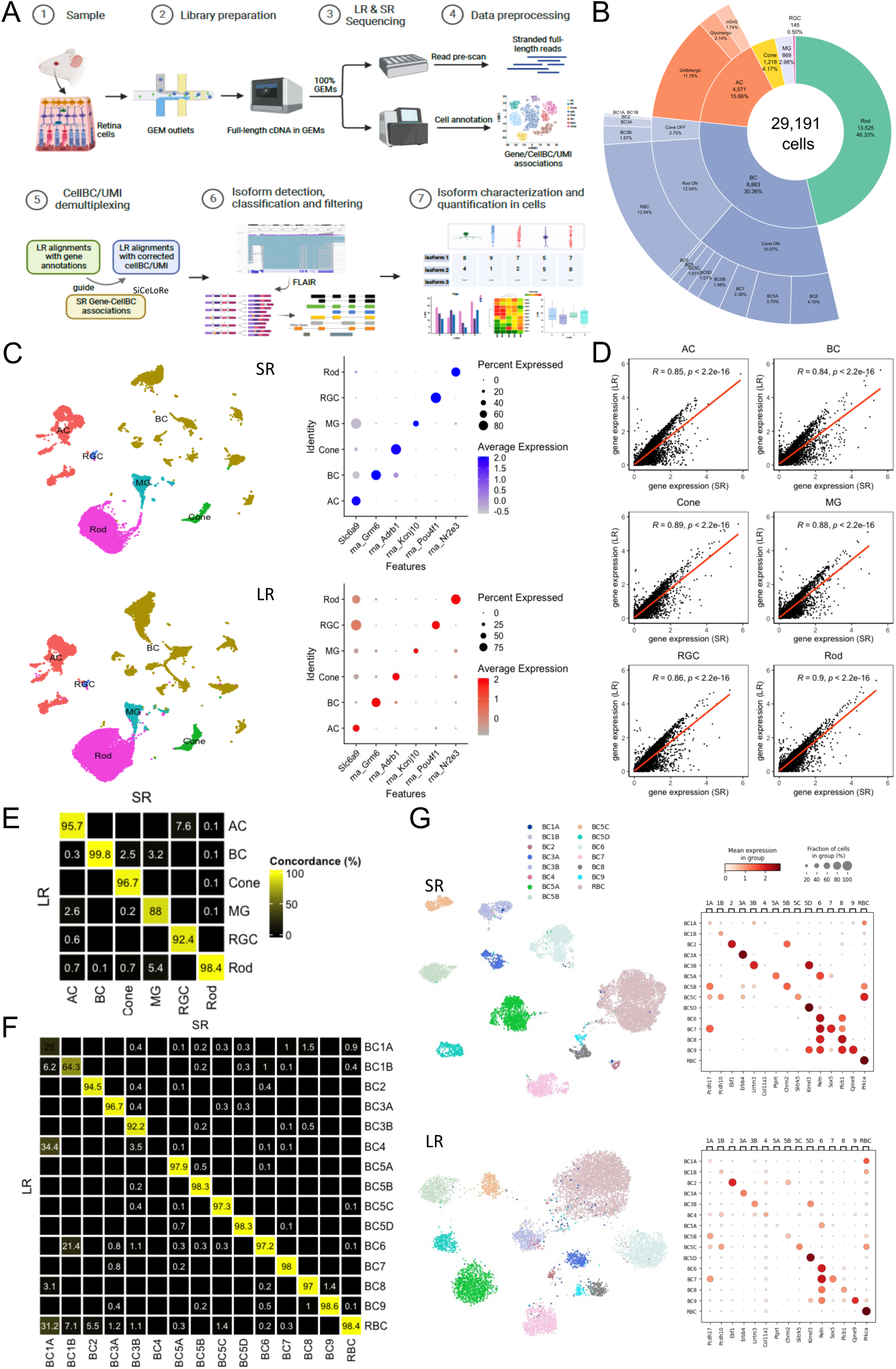
Overview of the experimental design and basic summary statistics between Illumina short-read and ONT long-read scRNA-seq. (A) Summary of the study design, with an overview of the Illumina short read (SR) and ONT long read (LR) scRNA-seq data processing pipeline. (B) Summary of cell count in each major retina cell class and subclass. (C) UMAP visualization of cells in each major retina cell class for SR (left upper) and LR (left lower) and dot plots for viewing the expression level of known major cell class markers between SR (right upper) and LR (right lower) data. (D) Scatter plot for single-cell gene expression levels calculated by two approaches in each major retina cell class. (E) Concordance of major cell class annotation results between SR and LR. Only cells detected in both approaches were shown here. (F) Concordance of BC type annotation results between SR and LR approaches. Only cells detected in both approaches were shown here. (G) UMAP visualization of cells in each BC type for SR (left upper) and LR (left lower) and dot plots for viewing the expression level of known BC type markers between SR (right upper) and LR (right lower) data.

## Results

### Single cell RNA-seq with short- and long-read technologies

To benchmark and assess the performance of short read and long read single-cell sequencing technologies comprehensively, we performed the two technologies on cells isolated from four mouse retina samples. These included two biological replicates from wild-type retinas and two samples enriched in amacrine (AC) and bipolar cells (BC), leading to a more comprehensive coverage of rare cell types. A thorough comparison of short-read and long-read sequencing methods was conducted, as detailed in **Supplementary Table 1**.

In total, the transcriptomes of over 30,000 single cells from four mouse retinas were profiled, generating 1.54 billion Illumina short reads and 1.40 billion Oxford Nanopore long reads. The sequencing achieved high read coverage, with an average of over 45,000 reads per cell, and exhibited comparable depth between the short- and long-read datasets. The median read length for Oxford Nanopore Technologies (ONT) was about 1,000 base pairs, corresponding to the average size of full-length transcripts. The average base call quality score was approximately 12.5, corresponding to an estimated error rate of around 5.6% (**Supplemental Figure 1**).

Cell clustering and annotation were initially conducted on the short-read scRNA-seq dataset. Following the analysis pipeline previously described (Li et al. 2024), a total of 29,191 cells that passed quality filters were integrated and clustered, forming six major groups (**Fig. 1B** and **1C**). Cell clusters were annotated using established cell class-specific marker genes (Li et al. 2024) (**Supplemental Figure 2**). All major cell classes in the retina were identified, including 13,525 rod cells, 8,863 BCs, 4,571 ACs, 1,218 cone cells, 869 Müller glial cells (MG), and 145 retinal ganglion cells (RGC). As expected, the two biological replicates of wild-type retina displayed a similar distribution of cell type proportions, with rod cells being the most abundant (**Supplemental Table 2**). In contrast, the samples enriched for ACs and BCs showed a significant increase in the proportion of the corresponding cell classes (**Supplemental Table 2**). Given the known heterogeneity within AC, BC, and RGC classes, based on their distinct transcriptomic, morphological, and functional properties, notable cell heterogeneity was observed, particularly in AC and BC clusters. To attain higher resolution, ACs were further clustered into four subclasses: GABAergic, Glycinergic, and non-GABAergic non-Glycinergic (nGnG, also recognized as starburst AC), while BCs were further clustered into 14 cell types (**Fig. 1B**).

To compare the performance of long-read to short-read sequencing, we conducted cell clustering and annotation using long-read data generated from the same set of samples. As depicted in **Fig. 1A**, long reads that passed quality filtering were demultiplexed based on cell barcodes (CB) identified by short-read sequencing (details in Methods), resulting in approximately 518 million long reads across the four samples (**Supplemental Table 3**). Upon mapping, cell clustering was performed, and the resulting clusters were annotated using known marker genes, following the same pipeline as the short-read dataset. Consistently, six major cell classes were identified with cell proportions similar to the short-read approach (**Fig. 1C**, **Fig. 1D** and **Supplemental Figure 2**). The consistency of cell class annotations between the short-read and long-read datasets was analyzed by comparing individual cell annotations (**Fig. 1E**). Remarkably, over 98.0% of cell class assignments (28,606 out of 29,191 cells) were consistent, with BCs showing an agreement of 99.8%. This high level of concordance was also evident at higher cell cluster resolutions. For instance, in BC cell types, a concordance of 97.4% (8,629 out of 8,863 cells) was observed at the individual cell type level (**Fig. 1F**). Interestingly, the long-read data identified an additional BC type, BC4, that was missed in the short-read data (**Fig. 1G**).

We further assessed the correlation of gene expression by comparing the transcriptomics across all cell classes from both datasets. As illustrated in **Supplementary Figure 3**, a strong positive correlation in gene expression was observed, with an overall Pearson’s r value of 0.87 (p-value < 2.2e−16). This strong correlation was consistent across all cell classes, with Pearson’s r values ranging from 0.84 to 0.90 (**Fig**. **1D**, p-value < 2.2e−16). Therefore, our results indicated that when sequenced at similar depths, both short-read and long-read datasets exhibited comparable sensitivity and high concordance in cell identification, clustering, and annotation.

### Isoform catalog in mouse retina and the unique splicing profiles across cell classes

One of the key advantages of long read is the improved ability of detecting transcript isoforms. To assess the robustness of the long-read sequencing approach in discovering splicing isoforms of different cell classes in mouse retina, we performed a detailed isoform analysis including identification, classification and quantification on the long-read data.

Under a stringent cutoff, as shown in **Fig. 2A**, using annotated transcripts in GENCODE (vM25) as the reference, a total of 44,325 transcript isoforms were identified. Most transcript isoforms (38,247, 86.2%) belonged to protein-coding genes. About 60% of isoforms match with known isoforms, including 19,821 isoforms matching a reference transcript at all splice junctions (Full Splice Match, FSM) and 7,517 isoforms partially matching to consecutive splice junctions of the reference transcripts (Incomplete Splice Match, ISM). Strikingly, the remaining 40% represents novel isoforms, including 15,894 isoforms of known genes with novel splice junctions of known splicing sites (Novel in catalog, NIC), 38 isoforms of known genes with novel donor and acceptor sites (Novel not in catalog, NNC), and 1,055 isoforms matching the concatenation of two or more separate genes (Fusion). It was interesting to note that novel isoforms (NNC, NIC and fusion) tended to be expressed at lower levels (**Fig. 2B**), which could provide a partial explanation for why these isoforms remained undetected in previous studies. Therefore, single-cell long-read sequencing greatly increased the number of isoforms detected.

**Figure 2.**
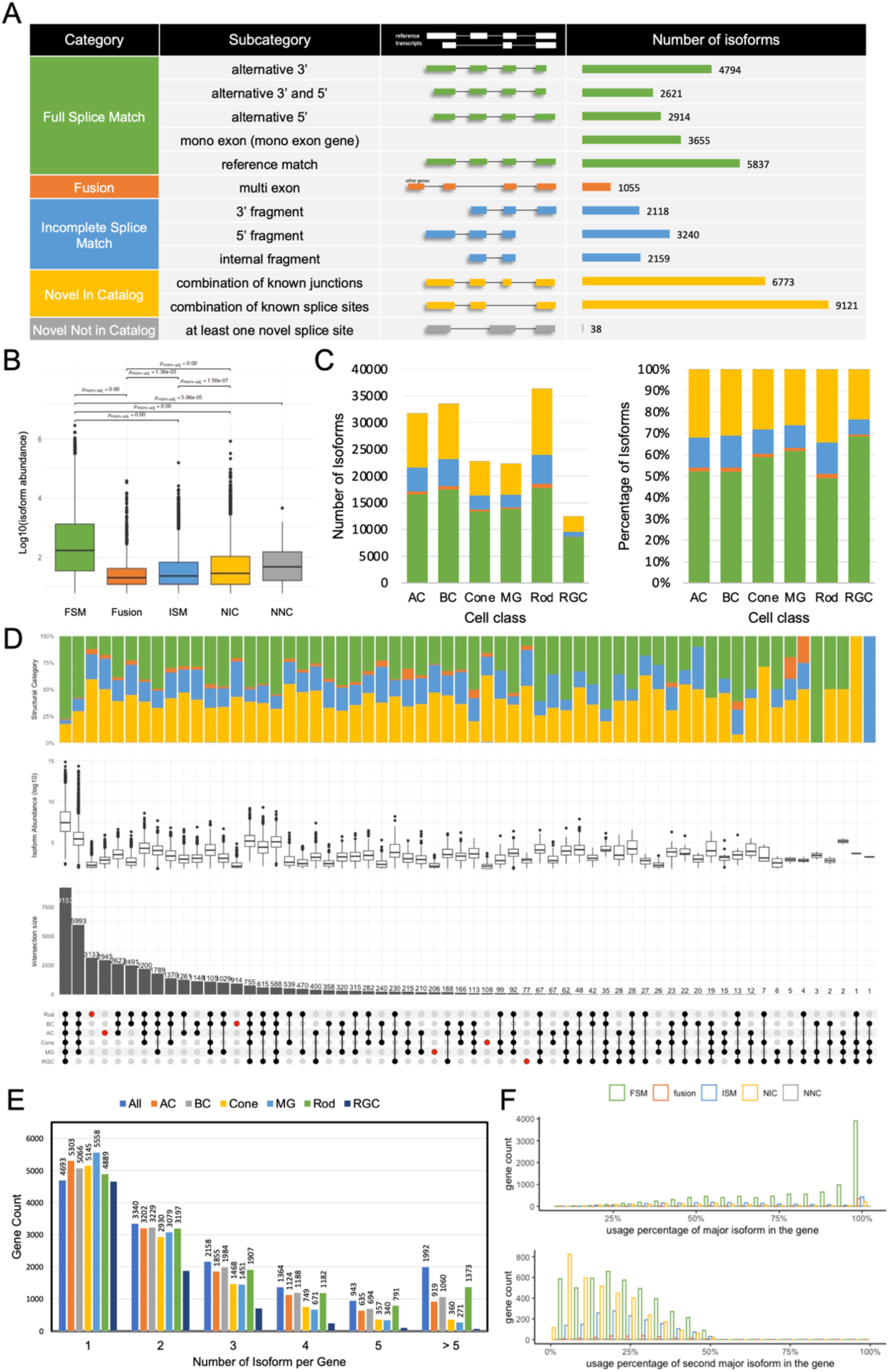
Overview of single-cell isoform catalog in the mouse retina. (A) Classification of isoforms according to their splice sites when compared to reference annotations and the number of isoforms in each category detected by the entire dataset. (B) Box plot showing the isoform abundance in different categories specified in (A). The adjusted p-value shows the group differences. (C) Summary of number (left) and percentage (right) of isoforms in different categories for each cell class, colored by categories specified in (A). (D) UpSet plot showing overlap of isoforms in cell classes, where number and percentage of isoforms shared by different cell classes were indicated in the top bar charts, colored by categories specified in (A), and the isoform abundance in each group were showed in the box plot. (E) Bar plot of the number of distinct isoforms expressed per gene. Genes with more than five distinct transcripts were merged. (F) Histogram showing the differential transcript usage in gene of the two most abundant isoforms of each gene, grouped and colored by categories specified in (A).

The number of novel isoforms identified across different cell classes varied significantly, with over 13,000 novel isoforms found in rod cells followed by BC and AC (**Fig. 2C**), correlating with the number of cells from each cell class profiled in our dataset (**Fig. 1B**). Despite the difference in raw isoform numbers, overall similar distribution of different isoform categories across cell classes was observed with known isoforms accounting around 65% (**Fig. 2C**). In contrast, the proportion of different isoform categories varied significantly depending on their expression pattern (**Fig. 2D**). For example, out of the transcript isoforms that were expressed across all cell classes, approximately 20% of these isoforms were newly discovered. In contrast, for isoforms that showed a more restricted expression pattern, the proportion of novel isoforms was generally higher. It was worth noting that although the vast majority of isoforms were expressed in at least three cell types, about 16.7% of isoforms were expressed exclusively in one cell class, including 2,945, 914, 108, 206, 77, and 3,133 isoforms identified for AC, BC, cone, MG, RGC, and rod, respectively.

Consistent with previous studies (Tian et al. 2021), an average of four isoforms, ranging from 2 to 28, were observed for the majority of genes (68%) (**Fig. 2E**). Similar distribution was observed across all cell classes (**Fig. 2E**). Subsequently, we classified the structural categories (**Supplemental Table 4**) and compared the expression levels (**Fig. 2F**) of the two most abundant isoforms for each gene. Strikingly, approximately 34% of genes exhibited a novel isoform among their top two isoforms. Moreover, for about half of the genes (6,336 out of 14,490) expressed in the retina, the major isoform constituted over 90% of the total gene expression, which indicated these genes had a dominant isoform.

Several features of known and novel isoforms were compared, including the number of exons (**Fig. 3A**), differential transcript usage within genes (**Fig. 3B**), and the length of the open reading frame (ORF) (**Fig. 3C**). The results revealed that while the distribution of exon numbers and ORF lengths between known and novel isoforms exhibited similar patterns, their usage within genes exhibited significant differences. Most of the known isoforms accounted for over 90% of the total gene expression, whereas the novel isoforms showed lower expression levels. We then extended our comparison to examine the differential usage within genes of known and novel isoforms across different cell classes (**Fig. 3D**), and they exhibited consistent trends. This pattern also held true for isoforms specific to cell classes (**Fig. 3E**).

**Figure 3.**
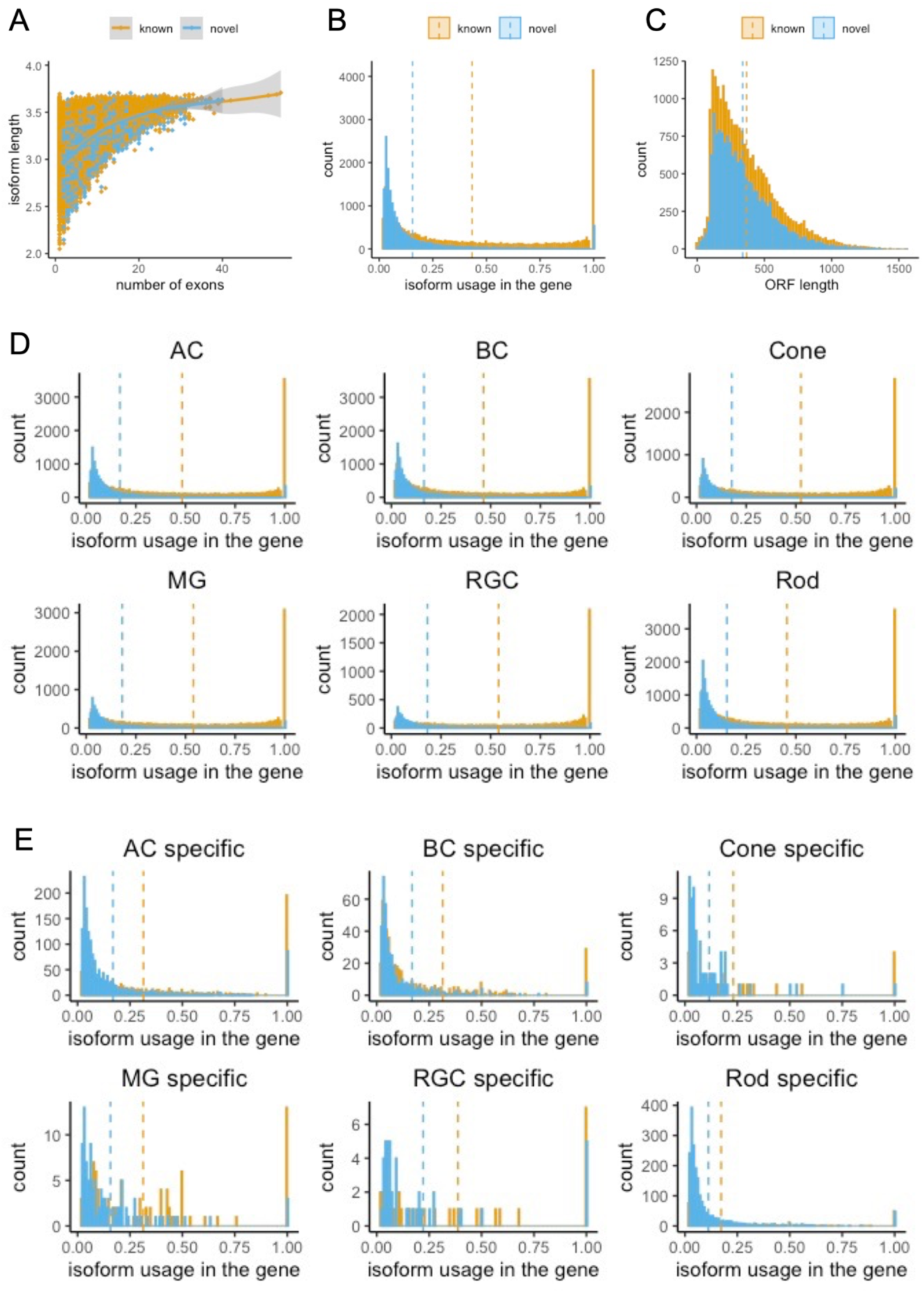
Comparison of novel and known isoforms in general and across different cell classes. (A) Scatter plot showing isoform length and exon numbers for known and novel isoforms. (B) Histogram showing the distribution of known and novel isoforms on expression level in the gene. (C) Histogram showing the distribution of known and novel isoforms on open reading frame (ORF) length. (D) Histogram showing the distribution of known and novel isoforms expressed in different cell classes on expression level in the gene. (E) Histogram showing the distribution of cell class-specific known and novel isoforms on expression level in the gene.

### Most genes displaying varied isoform usage among cell classes/subclasses/types

Since the vast majority of genes expressed multiple isoforms, it was interesting to examine whether genes demonstrated differential transcript usage (DTU) among the cell classes, subclasses or types illustrated in **Fig. 1B**. Considering the high dropout rate in single-cell data, DTU analysis was performed by merging cells from the same cluster into a pseudo-bulk sample to calculate the proportion of isoform usage across different cell classes. Consistent with the observation where only a small portion of isoforms showed cell class specific expression, most if not all isoforms of a given gene tended to be expressed across all cell classes. However, the proportion of different isoforms varied significantly among cell classes. For example, although all isoforms of *Pcbp4* were expressed in all cell classes, varied usage among different cell classes was observed (**Fig. 4A** and **4B**). As shown in **Fig. 4B**, the transcript (colored in yellow) that was predominantly expressed in ACs was lowly expressed in rods, exhibiting significant difference (p-value = 1.06E-88, **Fig. 4C**). Conversely, a transcript (colored in dark blue) was significantly more prevalent in rods than in BCs (p-value = 3.86E-98, **Fig. 4C**). Similarly, for the two isoforms of *Prkcz*, the isoform comprising 15 exons was predominantly expressed in ACs, BCs, and RGCs, while the isoform with 18 exons accounted for over 90% expression in cones, MGs, and rods (**Fig. 4D**).

**Figure 4.**
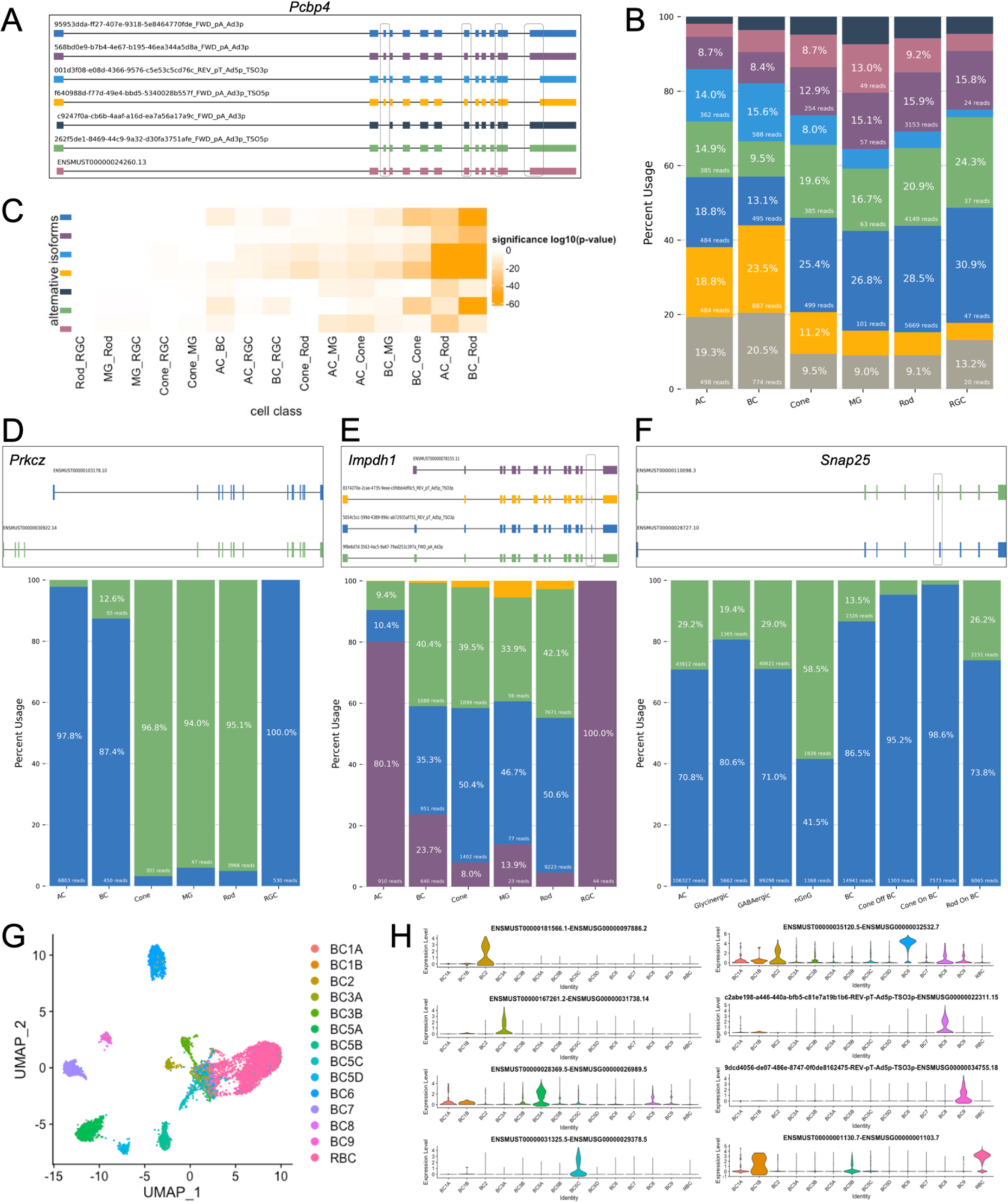
Statistics of differential usage of transcript isoforms across cell classes and subclasses. (A) Top 7 most abundant transcript isoforms of gene *Pcbp4* detected. Differentially spliced sites were outlined in black. (B) Percent usage of each *Pcbp4* isoform, with colors corresponding to isoforms in (A). All other less abundant isoforms not plotted in (A) were merged and represented as grey bars. (C) Heatmap showing the significance of alternative isoform (colored by isoform specified in (A)) usage between two major cell classes using Fisher’s exact tests. The greater the disparity in isoform usage between the first and second cell classes, the lower the p-value. (D) Isoform plots and usage bar charts of *Prkcz* showing different isoform expression patterns across major retina cell classes. (E) Isoform plots and usage bar charts of *Impdh1* showing different isoform expression patterns across major retina cell classes. (F) Isoform plots and usage bar charts of *Snap25* showing different isoform expression patterns across amacrine and bipolar cell subclasses. (G) UMAP visualization of 8,863 BCs clustered using isoform as features and colored by BC subclass annotations from SR scRNA-seq. (H) Violin plot showing selected isoforms differentially expressed across BC subclasses.

Another example is *Impdh1*, which has associations with inherited human retinal disorders like Leber congenital amaurosis (LCA) and retinitis pigmentosa (RP). We identified several novel isoforms of *Impdh1*, which include a novel 17bp exon previously unreported in the Reference Sequence (RefSeq) transcripts (**Fig. 4E** upper). The inclusion of the 17bp exon resulted in a reading frameshift and ORF elongation (37 amino acids) with an alternative stop codon. The transcripts incorporating this exon demonstrated significant expression in BCs, cones, MGs, and rods (**Fig. 4E** lower). The primary isoform of *Impdh1* expressed in mouse photoreceptors was not the canonical *Impdh1*, suggesting that changes in the canonical protein’s function were probably not the primary cause of retinal degeneration. Additionally, this exon exhibited a high degree of conservation in humans, and we identified several single nucleotide variants located within 10 base pairs upstream or downstream of the novel exon in our in-house RP patient cohort, as indicated in **Supplemental Table 5**. These variants were not present in the general populations in the Genome Aggregation Database (gnomAD) (Karczewski et al. 2020). It would be intriguing to explore its correlation with human diseases in subsequent studies.

Subsequently, we analyzed the pattern of isoform usage within the subclasses of amacrine and bipolar cells. Mirroring what we observed in major cell classes, it appeared that most isoforms were present across all subclasses, though the distribution of these isoforms shifted between subclasses. For example, as shown in **Fig. 4F**, one isoform of *Snap25* (colored in blue) was mainly found in GABAergic and Glycinergic subclasses, while another one (colored in green) constituted over half of the expression in nGnG ACs. Furthermore, the expression ratios of these two isoforms varied between cone BCs and rod BCs.

To assess transcript isoform usage in individual cell types, we further counted the isoform usage for each individual bipolar cell, resulting in a cell-by-isoform matrix. We then undertook cell clustering with isoforms serving as distinguishing features (**Fig. 4G**). While no additional cell clusters emerged compared to gene expression-based annotations, we observed differential expression patterns of isoforms among BC types and identified several isoforms that were predominantly expressed in specific BC types (**Fig. 4H**). Notably, some of these were novel isoforms.

In summary, our study revealed that genes exhibited diverse isoform usage patterns across major retina cell classes and subclasses. Additionally, we have identified distinct expression patterns of isoforms across BC types, as well as isoforms that are primarily expressed in specific cell types.

### Gene fusion isoforms in mouse retina

A surprising finding was the identification of a substantial number (1,055) of potential fusion transcripts. These fusion transcripts were unlikely due to experimental artifacts as supported by the following evidence (1) The fusions were substantiated by a minimum of 5 UMIs from long reads that spanned a fusion breakpoint (details in Methods), therefore indicating these were independent events and unlikely to be artifact. (2) All detected fusions occurred within the same chromosome, with no indication of inter-chromosomal fusions, indicating it was unlikely due to template switch error during polymerase chain reaction (PCR). (3) Notably, the genes involved in each fusion were found on the same strand, further suggesting that these fusion transcripts were biologically relevant events.

Most of these fusions were characterized by the combination of two genes, though instances of three-gene fusions were observed (**Fig. 5A** and **5C**). Subsequently, we assessed the proximity of the fused genes. Some exhibited adjacent fusion, while others skipped over nearby genes to fuse with more distant counterparts (**Fig. 5B** and **5C**). When examining the exon count and abundance of fusions, categorized by gene proximity, it became evident that adjacent fusions contained a greater number of exons compared to the non-adjacent counterparts (**Fig. 5D**, adjusted p-value = 1.8E-07), and variations in abundance were noted between the two groups (**Fig. 5E**, adjusted p-value = 0.0073).

**Figure 5.**
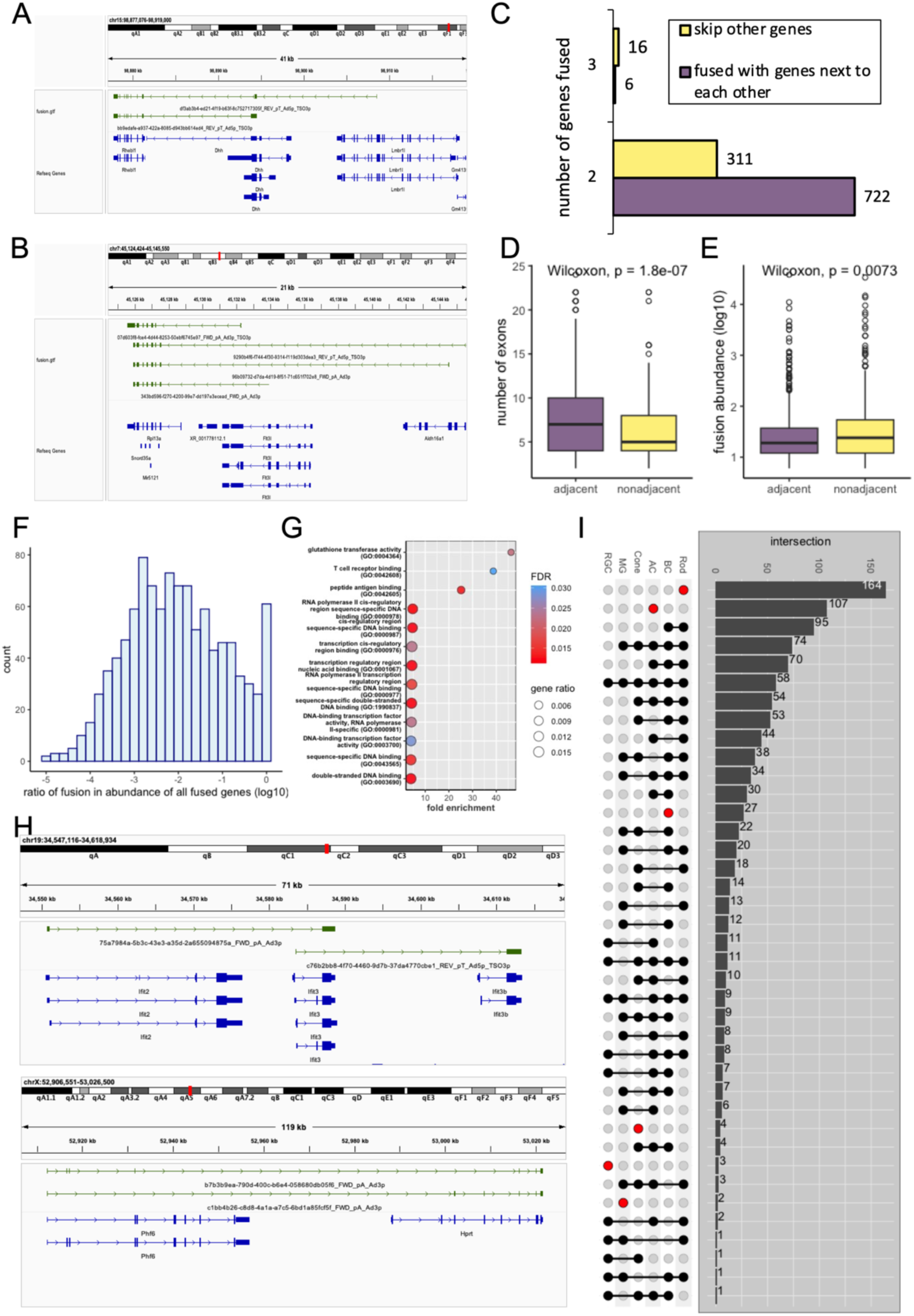
Gene fusions in mouse retina. (A) Example visual of fusions (in green) aligned to the RefSeq reference (in blue) formed by 2 or 3 known genes. (B) Example visual of fusions (in green) aligned to the RefSeq reference (in blue) formed by adjacent or distant known genes. (C) Bar chart illustrating the count of fusions, categorized by the number of fused genes and their adjacency to one another. (D) Boxplot showing the exon number of fusions categorized by the adjacency of the fused genes. (E) Boxplot showing isoform abundance of fusions categorized by the adjacency of the fused genes. (F) Histogram illustrating the distribution of fusion expression ratios compared to the combined expression of fused genes, with log transformation applied. (G) Bubble chart depicting the 13 molecular functions of Gene Ontology enrichment analysis ranked by fold enrichment, utilizing gene set always in fusions. (H) Example visuals of fusions (in green) aligned to the RefSeq reference (in blue). The top illustration depicted a gene (*Ifit3*) that can fuse with multiple other genes (*Ifit2* and *Ifit3b*). The lower visualization highlighted the alternative splicing within the fusion (*Phf6*-*Hprt*). (I) UpSet plot showing intersection of fusions in major retina cell classes, where number of fusions shared by different cell classes were indicated in the right bar charts.

We then compared the abundance of the fusions against the expression of all isoforms present in the implicated genes. The proportions between fusion utilization and other isoform usage showed a broad distribution (**Fig. 5F**). Intriguingly, 90 genes were exclusively expressed in fusions. Functional clustering of these genes unveiled common pathways (**Fig. 5G**), notably those associated with immunity, such as the T cell receptor binding (GO:0042608), which was logical considering the need for adaptability in immune processes.

In addition, upon detailed examination of the detected gene fusions, we also observed that: (1) Certain genes could partner with multiple other genes, resulting in different fusions (**Fig. 5H**, upper). (2) Some fusions underwent alternative splicing events (**Fig. 5H**, lower), which would be interesting for cancer and drug target related studies. We further evaluated the overlap of fusions across various cell classes (**Fig. 5I**) and pinpointed certain fusions unique to each cell class, tallying 164 in rods, 107 in ACs, and 27 in BCs.

### Down-sampling analysis and sequencing saturation

To evaluate the sensitivity of sequencing depth on isoform detection and determine if our sequencing reached saturation, we conducted simulation by randomly sampling 1%, 10%, and 50% of the dataset and performed the analysis with the same pipeline. As expected, the number of isoforms detected positively correlates with the number of sequencing reads used in the analysis (**Fig. 6A**). However, the increasing rate varied among different structural categories. For example, the number of FSMs detected increased by 41.3% from 10% to 50% of the full dataset and only increased slightly by 10.2% from 50% to the full dataset (**Fig. 6A** and **Supplemental Figure 4A**), indicating they were approaching saturation and canonical isoforms can be detected by a moderate number of reads. Conversely, the quantity of novel isoforms, such as NICs, kept escalating. There was a 111.3% increase in their numbers when comparing 10% of the full dataset to 50%, and a further 26.4% increase was observed when expanding from 50% of the dataset to the entire dataset, indicating that it has not been saturated even with the full dataset, probably due to their low expression level (**Fig. 6A** and **Supplemental Figure 4A**).

**Figure 6.**
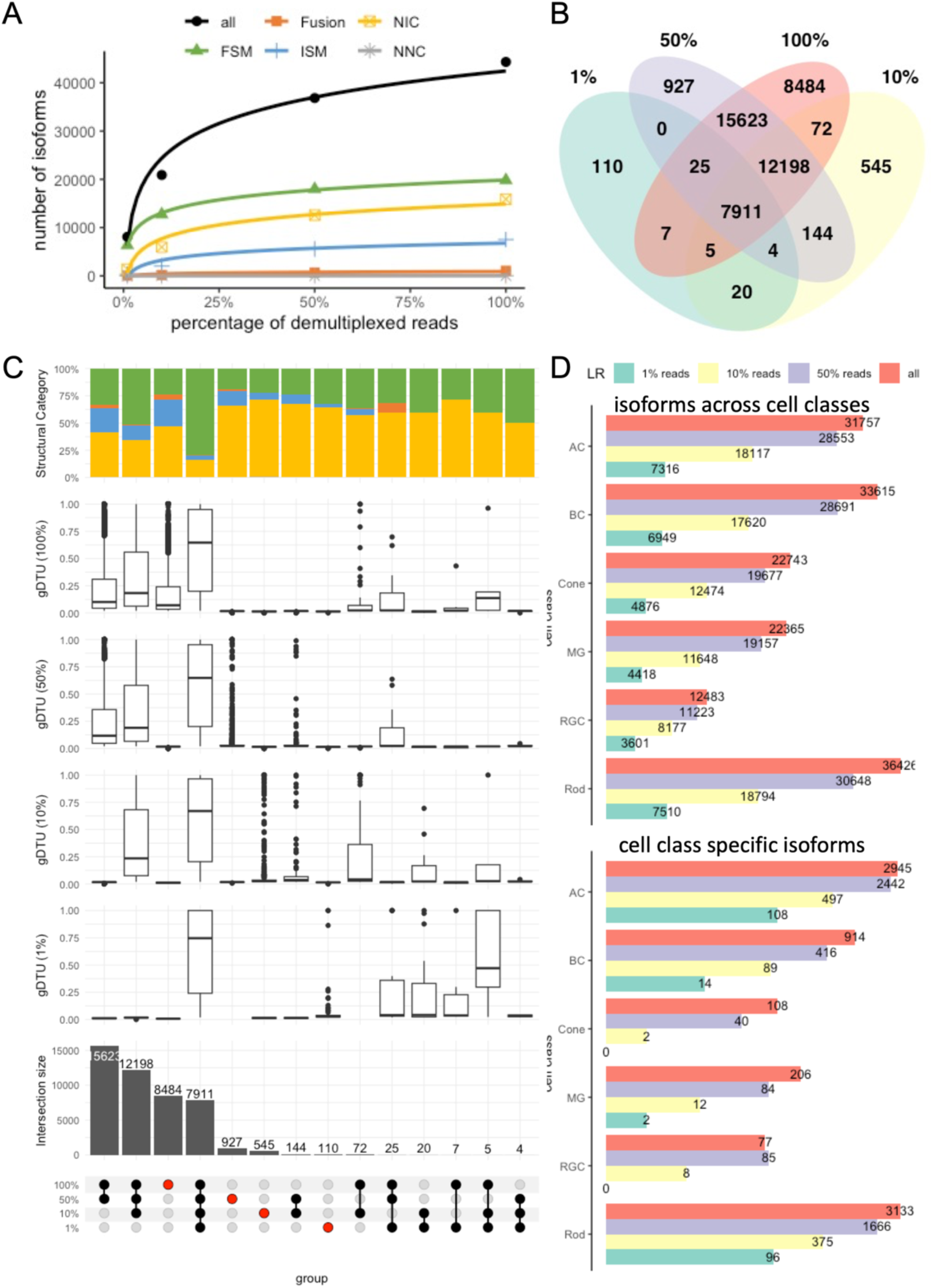
Down-sampling analysis. (A) Classification of isoforms according to their splice sites when compared to reference annotations and number of isoforms in each category detected using 100%, 50%, 10% and 1% of demultiplexed LRs. (B) Venn diagram showing intersection of the isoform sets detected using all, 50%, 10% and 1% of demultiplexed LRs. (C) UpSet plot showing intersection of isoforms detected using 100%, 50%, 10% and 1% of demultiplexed LRs, where number and percentage of isoforms shared by different cell classes are indicated in the bottom and top bar charts, colored by categories specified in (A). The box plot shows the DTU in gene before filtering with 2% cutoff for the isoforms in each group. (D) Bar plot shows number of all isoforms (above) and cell class specific isoforms (bottom) in each major retina cell class detected using 100%, 50%, 10% and 1% of demultiplexed LRs.

We further examined the overlap of isoforms detected using 1%, 10%, 50%, and all long reads. Interestingly, some isoforms were identified in datasets with fewer sequencing reads but were absent when more reads were used (**Fig. 6B**). For instance, 1,075 isoforms were detected using 50% of the long reads, but they were not present in the comprehensive isoform set derived from all long reads. This discrepancy arose because we excluded isoforms that had a usage of less than 2% of the total usage for their respective genes, in order to mitigate background noise (**Fig. 6C**). Thus, when we down sampled the reads, the isoforms with low abundance could be kept by chance if their proportion becomes higher than 2%.

As we increased the number of reads utilized for isoform identification, the number of isoforms detected in each cell class followed a similar upward trend (**Fig. 6D**, upper) and the fraction of FSMs within the identified isoform set saw a downtick (**Supplemental Figure 4C-E**). The count of identified isoforms in the six retina cell classes experienced a notable rise, ranging from 30% to 60%, when comparing the data from 10% of the full dataset to 50%. Subsequently, when expanding from 50% of the dataset to the complete dataset, a further increase ranging from 11% to 18% was observed. This pattern suggested that isoform detection within each cell class approached a state of saturation. In addition, upon examining the overlap of isoforms across cell classes for each dataset, we observed that 21.75% (8,012 out of 36,832), 26.58% (5,555 out of 20,899), and 27.32% (2,208 out of 8,082) isoforms from sets determined using 50%, 10%, and 1% of long reads, respectively, were expressed across all cell classes. This compared to 20.65% (9,153 out of 44,325) in the complete dataset, suggesting a smaller number of reads more tend to identify the common isoforms across cell classes. Additionally, the identification of cell class-specific isoforms has not yet reached saturation, as evidenced by a significant 56.0% increase in isoform number when the dataset was expanded from 50% to its full extent (**Fig. 6D**, lower).

## Discussion

Short-read scRNA-seq, now as a standard method, is highly effective in profiling gene expression to identify cell types and trajectories (Hwang et al. 2018), however, there’s a limitation in the length of the sequenced reads. Short-read scRNA-seq can be divided into two main approaches: transcript counting methods (e.g., 10x Genomics) that sequence transcript ends, and whole transcript methods (e.g., Smart-Seq2 (Picelli et al. 2013)) that cover entire RNA sequences. Transcript counting methods can profile many cells but have limited capacity for splicing or isoform information, whereas whole transcript methods provide more details on exon alternative splicing but offer lower throughput. Previous short-read scRNA-seq studies employing whole transcript methods have revealed significant variations in isoform expression among individual cells (Shalek et al. 2013; Marinov et al. 2014). However, these analyses predominantly concentrated on alterations in specific exons or splice junctions, a limitation imposed by the nature of short-read sequencing. The reconstruction of isoforms remained suboptimal for sequences longer than 1 kb, with only about 40% of molecules assignable to a specific isoform (Hagemann-Jensen et al. 2020). This technological limitation leaves a gap in understanding the full extent of alternative splicing and the diversity of isoform expression both within and across single cells and underscores the need for improved methodologies in accurately capturing and characterizing longer isoforms.

Long-read sequencing technology aims to resolve this issue entirely; however, its adoption in single-cell research is currently hindered by the lack of established protocols and data analysis pipelines (De Paoli-Iseppi et al. 2021). In our study, we developed a workflow designed to streamline the analysis aspect of high-throughput long-read RNA sequencing data, enabling the identification of transcript isoforms at the single-cell level. Our workflow modified the 10x Genomics scRNA-seq protocol (Gupta et al. 2018) in order to comprehensively resolve the full-length transcriptome of the mouse retina, and this approach can be applied to other cDNA libraries. In comparing the long-read and short-read datasets, we discovered highly comparably results in both gene expression and cell annotation. Our analysis revealed a 98.0% consistency in cell class categorization between long-read and short-read datasets, and a closely aligned 97.4% agreement at the bipolar cell type level. In short, the long-read sequencing approach demonstrated its reliability in detecting single-cell transcriptomes based on gene expressions, making it plausible to use long read sequencing exclusively for scRNA-seq in the future (Shiau et al. 2023).

The integration of long-read sequencing with single-cell sequencing techniques holds the promise of filling the existing gaps in isoform information. Using this pipeline, we conducted the first comprehensive characterization of full-length transcription isoforms within individual cells of the mouse retina. Over 16,000 novel transcripts were identified at lower expression levels compared to previously known ones. Additionally, we pinpointed 7,383 isoforms specific to cell classes, many of which were novel and exhibited low expression levels. The identification of cell-class-specific isoforms holds potential applications in various fields, such as immunotherapy, where cell surface proteins play a pivotal role. Interestingly, our analysis of the mouse retina showed the distribution of novel transcripts was consistent across different retinal cell classes and unveiled a common pattern where all major cell classes in the tissue expressed a combination of diverse isoforms rather than a single canonical isoform. We frequently observed intricate splicing variations between the two most abundant isoforms of a gene. Furthermore, many genes displayed varying patterns of isoform usage among different cell classes and subclasses. Another noteworthy finding was that retinal cells could express different isoforms, even when their overall gene expression levels did not significantly differ from each other. This implied that identifying cell-class-specific genes based solely on gene expression levels was not sufficient to characterize the vast transcriptional diversity. By examining the transcriptomic profiles of individual cells, researchers can gain a deeper understanding of how different isoforms are utilized in specific cellular contexts, shedding light on the intricacies of gene regulation and function within heterogeneous cell populations.

In addition to what we have demonstrated in this study, this dataset has a wide range of applications across various genomic research areas. One such application is the exploration of single-cell gene fusions, which can provide valuable information about aberrant gene interactions and potential drivers of diseases. By leveraging long-read scRNA-seq data, we were able to identify a total of 1,055 intra-chromosomal gene fusions within the mouse retina. These gene fusions exhibited intriguing characteristics. For instance, certain genes had the capability to partner with multiple other genes, resulting in diverse fusion events. Furthermore, some of these fusions underwent alternative splicing events, further diversifying the transcriptomic landscape. What is particularly noteworthy was that certain genes can fuse with others that are not immediately adjacent to them on the same chromosome, highlighting the complexity and flexibility of gene fusion events in the context of single-cell RNA sequencing data. These findings may have implications in various fields, including cancer research, where gene fusions and differential isoform expression can have significant clinical relevance.

In conclusion, the long-read scRNA-seq approach was proved to be highly effective in identifying both known and novel isoforms. Our study stands as the first comprehensive characterization of full-length transcription isoforms in single cells within the mouse retina. Our analytical approaches served as an initial foundation to tackle these inquiries, which can be applied to human samples, propelling isoform-focused research in single cell of aging and disease context.

## Methods

### Sampling and animal procedures

All mice were handled in accordance with the policies on the treatment of laboratory animals by the Institutional Animal Care and Use Committee (IACUC) at Baylor College of Medicine, and the studies were conducted in adherence to the Animal Models of Retinal Development and Diseases protocol approved by Biomedical Research and Assurance Information Network. Stock C57BL/6J mice were purchased from the Jackson Laboratories, Bar Harbor, ME (stock number: 000664). The C57BL/6J mice were maintained on a 12-hour light/dark cycle at 23°C, standard mouse LabDiet 5V5R (Purina, St. Louis, MO), and water provided ad libitum throughout the study. The mice were managed and housed by the Baylor College of Medicine Center for Comparative Medicine. Adult mice were euthanized using CO2 gas and isoflurane asphyxiation for 5 minutes, followed by cervical dislocation.

### Single cell cDNA library preparation and sequencing

Samples and scRNA-seq cDNA and library generation were described in paper “Comprehensive single-cell atlas of the mouse retina” (Li et al. 2024). Briefly, scRNA-seq cDNA and library was generated using 10x Genomics Chromium Single cell 3’ Reagents Kits v2 and v3, following the manufacturer’s instructions. Short-read sequencing library was sequenced on an Illumina Novaseq 6000 sequencer (151bases + 151bases).

### Nanopore long reads library preparation and sequencing

Nanopore library was generated from scRNA-seq cDNA following the Nanopore Single-cell transcriptomics with cDNA prepared using 10x Genomics protocol (version Jan2022) using the SQK-PCS111 Ligation Sequencing Kit. 35-55 fmol of the library was sequenced on PromethION FLO-PRO002 R9.4.1 flow cells. The following options were used: 72hr of run time, and real-time basecalling with High-accuracy mode.

### Illumina short-read data analysis

All four samples’ fastq data were processed using Cell Ranger (7.0.1) to create a gene count matrix for each. Contamination from background transcripts in the preserved true cells was mitigated using SoupX (Young and Behjati 2020). Subsequently, DoubletFinder (McGinnis et al. 2019) was employed to estimate and remove potential doublets, particularly focusing on those with a high proportion of simulated artificial doublets. The gene count matrices were then fed into the standard Seurat (Hao et al. 2021) pipeline, with SCTransform v2 undertaking the normalization process. The samples were clustered at a resolution of 0.6. Annotations for the retina cell class were added manually using well-known retina marker genes. The subclass annotation for AC and BC was obtained through the utilization of “single-cell ANnotation using Variational Inference” (scANVI)(Gayoso et al. 2022), using an in-house meta reference described in paper “Unified comprehensive single-cell atlas of the mouse retina”. Data integration from all four datasets was carried out using Seurat. **Fig. 1C** and **1F** illustrate the results of the clustering and cell annotation.

### Oxford Nanopore long-read data preprocessing

Basecalling was carried out on the raw Oxford Nanopore Technology fast5 data using Guppy (version 6.1.5). The Nanopore fastq reads were inspected for adapter sequences and poly(A/T) sequences of 15 nucleotides or longer. We identified polyA (or T) sequences with a minimum of 75% adenine content within 170 nucleotides from both read ends, generating stranded (forward) reads via sicelore (Lebrigand et al. 2020) (https://github.com/ucagenomix/sicelore) utilizing the NanoporeReadScanner.jar module. The refined fastq data were then used to align the genome using Minimap2 (Li 2018) (v2.24, https://github.com/lh3/minimap2) with the parameter “-ax splice -uf --MD --sam-hit-only -- junc-bed”, referencing the GRCm38/mm10 genome, and annotated with Gencode vM25. Using sicelore’s Nanopore_BC_UMI_finder.jar module, we identified the location of the barcode sequence for each alignment by searching the flanking sequence to the cell barcode. The cell barcodes and gene correlation discovered from the Illumina short-read data served as a reference to find and trim in the long reads, with the dynamic edit distance adjusting the maximal edit distance according to the complexity of the search set. The sequences found after the cell barcode were treated as UMIs (Unique Molecular Identifiers) and were trimmed accordingly. The processed BAM was utilized for further analysis.

### Detection and quantification of isoforms

We utilized Flair (Tang et al. 2020) (https://flair.readthedocs.io/en/latest/index.html) to summarize the alignment for each read by grouping reads with similar splice junctions (<5bp) to get a raw isoform annotation. The final nanopore-specific reference isoform assembly is made by aligning raw reads to the first-pass assembly transcript sequence using minimap2 to identify isoforms with high confidence.

flair correct -q LR.bed -g mm10.fasta -f gencode.v25.mm10.annotation.gtf -o LR_corrected.bed

flair collapse -q LR_corrected.bed -g mm10.fasta -f gencode.v25.mm10.annotation.gtf -m

/path/to/minimap2 -sam /path/to/samtools -o LR -r demultiplexed_reads.fq --stringent

Pseudo-bulk samples were produced by merging isoform counts at the level of cell class, cell subclass, and individual cells. Cell class groupings originated from the Seurat clustering depicted in **Fig. 1C**, while AC/BC sub class groupings were derived from scANVI clustering presented in **Fig. 1G**. The “flair quantify” function in Flair was utilized to create a UMI count matrix for each tier, in which rows correspond to chosen transcripts and columns represent groups (cell classes/subclasses).

flair quantify -r reads_manifest.tsv -i LR.isoforms.fa -m /path/to/minimap2 -sam /path/to/samtools -o LR.flair.quantify --isoform_bed LR.isoforms.bed --sample_id_only --generate_map --tpm --stringent

### Isoform classification

SQANTI3 (Tardaguila et al. 2018) (version 5.1.2, https://github.com/ConesaLab/SQANTI3) was used to compare the transcripts identified to the reference with rules mode. The isoform classification was extracted from the SQANTI3 result and plotted in **Fig. 2A**. Results from comparisons across different cell classes were plotted using the R package ComplexUpset. We ranked transcript abundance for each gene that had multiple isoforms and obtained the alternative splicing events from the most expressed isoform and the second most expressed isoform.

### Isoform filtering

We implemented several filters to hone the isoforms according to specific parameters. The first parameter was intrapriming events: isoforms showing 12 or more adenines at the genomic level within the 20 base pairs downstream of the Transcription Terminating Site (TTS) were filtered out. RT-Switching was the next parameter: we removed isoforms that may have been affected by reverse transcription errors, resulting in the formation of new, non-canonical splice junctions. Isoforms linked to degraded RNA, as indicated by the retention of intronic sequences, were also considered for exclusion. Gene abundance and isoform abundance were also taken into account: we eliminated genes with a Unique Molecular Identifier (UMI) count of 10 or less, and isoforms with a UMI count of 5 or less. Moreover, we excluded isoforms with a UMI count of less than 2% of the total UMI count for the respective gene to account for background noise. The SQANTI3 was used to annotate and filter based on the first three parameters, while the last three were applied manually after isoform quantification. These filtering steps were used to refine the isoform dataset, ensuring a more precise and reliable set of isoforms for additional analysis.

### Differential isoform usage analysis

Utilizing the matrix detailing isoform expression by cell class/subclass, the transcript structures and the percentage usage visualizations in **Fig. 4** were constructed via the “plot_isoform_usage” function in Flair. The significance of differential isoform utilization across cell classes was determined using the “diff_iso_usage” function in Flair, which is based on Fisher’s exact tests. After scaling the matrix of cell-by-isoform expression, the outcomes were incorporated into UMAP visualizations, as depicted in **Fig. 4** and facilitated by Seurat.

### Down-sampling analysis

The down-sampling datasets were achieved by randomly subsampling the demultiplexed long reads (1%, 10%, 50%) using seqtk (version: 1.3-r115-dirty, https://github.com/lh3/seqtk) and re-running the pipeline with the same parameters and filtering. The gffcompare(Pertea and Pertea 2020) (0.12.6) program was used with the command “gffcompare -i input -o out -r gencode.vM25.annotation.gff3 -R” to compare transcript isoforms annotations obtained from the 3 down-sampling datasets and the complete isoform catalog. Results from these comparisons were plotted using R packages VennDiagram (**Fig. 6B**) and ComplexUpset (**Fig. 6C**).

## Data Access and Code Availability

All raw sequencing data, a gene transfer format (GTF) file including all transcript isoforms passed filtering and a cell class by isoform count matrix generated in this study have been submitted to the NCBI Gene Expression Omnibus (GEO; https://www.ncbi.nlm.nih.gov/geo/) under accession number GSE255520. Refer to Table S1 for a summary of these datasets. USCS genome browser tracks of the analyzed transcript isoform data are available at https://genome.ucsc.edu/s/wmeng1018/SC_transcript_isoform_mouseRetina. The code and scripts used in this study are available from https://github.com/RCHENLAB/LR_scRNA-seq_manuscript

## Acknowledgments

This work was supported by the Rui Chen Lab at the Baylor College of Medicine and was funded by Chan Zuckerberg Initiative [CZF2019-002425]. We would like to thank eyeGENE for providing patient samples collected at the National Eye Institute. We acknowledge Dr. Salma Ferdous for her contribution to improving the proofreading and English language in our manuscript.

## Author Contributions

M.W., R.C. and Y.L. conceived the study design. Y.L. collected mouse samples, performed experiments and sequencing. M.W. developed the pipeline, conducted data analysis and wrote the manuscript while J.W., and R.C provided feedback. S.O. provided helpful comments and discussion. R.C. planned and supervised the research and wrote the manuscript. All authors provided critical feedback, read and approved the final manuscript.

